# Dopaminergic signalling modulates reward-driven music memory consolidation

**DOI:** 10.1101/2020.04.01.020305

**Authors:** Laura Ferreri, Ernest Mas-Herrero, Gemma Cardona, Robert J. Zatorre, Rosa M. Antonijoan, Marta Valle, Jordi Riba, Pablo Ripollés, Antoni Rodriguez-Fornells

## Abstract

Previously, we provided *causal* evidence for a dopamine-dependent effect of intrinsic reward on memory during self-regulated learning (Ripollés et al., 2016; Ripollés et al., 2018). Here, we further investigated the dopamine-dependent link between reward and memory by focusing on one of the most iconic *abstract* rewards in humans: music. Twenty-nine healthy participants listened to unfamiliar excerpts—which had to be remembered following a consolidation period—after the intake of a dopaminergic antagonist, a dopaminergic precursor, and a placebo across three separated sessions. The intervention modulated the pleasantness experienced during music-listening and memory recognition of the presented songs (i.e., lower with the antagonist, higher with the precursor) in individuals with higher sensitivity to musical reward. Our work highlights the flexibility of the human dopaminergic system, which is able to enhance memory formation not only through explicit and/or primary reinforcers but also via intrinsic, abstract, or aesthetic rewards of different natures.

## Introduction

Reward lies at the center of human behavior and is hard-wired into the brains of all vertebrates, from humans to birds (Schultz, 2015). As such, much research—including extensive work in animal models—has been devoted to understanding its underlying neurobiological mechanisms (Berridge and Kringelback, 2008). Of the many cognitive processes related to reward, one has received particular attention in recent years: pleasure (i.e., liking or hedonic reward, Berridge and Kringelback, 2008). Pleasure is an especially complex construct in humans, who not only respond to primary (e.g., food) and secondary reinforcers (e.g., money, Delgado et al., 2003) but can also experience intrinsic rewards triggered by internal mental states (e.g., flow and curiosity, Csikszentmihalyi, 2014; Gruber et al., 2014) or intrinsic motivational processes (e.g., Berlyne, 1960; Ryan & Deci, 2000; Gottlieb et al., 2013). In addition, pleasure is intimately related to multiple cognitive processes such as learning and memory (Shohamy et al., 2010; Lisman et al., 2011; Redondo et al., 2011). In this vein, we recently showed: i) that humans, even in absence of explicit feedback, can experience pleasure from the process of learning itself (Ripollés et al., 2014, 2016); ii) that this *intrinsic* reward signal modulates the entrance of new information into long-term memory via the activation of the dopaminergic midbrain, hippocampus, and ventral striatum (the SN/VTA-Hippocampal loop, a brain circuit in the service of memory and learning; Lisman and Grace, 2005; Ripollés et al., 2016); and most importantly iii) that dopamine plays a *causal* role in this process, with its effects being further modulated by individual differences in general sensitivity to reward (i.e., the dopamine-dependent memory effects induced by intrinsic reward are greater in more hedonic participants; Ripollés et al., 2018).

Our previous work thus explored the neural mechanisms of intrinsic reward and its effects on memory in the context of self-regulated learning (i.e., the reward we experience from learning on our own, without external feedback). Humans experience pleasure from a great many other types of abstract pleasure, however (O’Doherty et al., 2001; Sescousse et al., 2013; Bhanji and Delgado, 2014). Among abstract reinforcers, *music* has received special attention in recent years as a powerful vector for studying pleasure. Music-induced pleasure relies on the activity of core reward regions within the mesolimbic *dopaminergic* system (Blood and Zatorre, 2001; Salimpoor et al., 2011, 2013; Koelsch, 2014; Mas-Herrero et al., 2018; Ferreri et al., 2019). Both theoretical considerations and recent experimental findings suggest that music represents a learning challenge by itself—triggered by the presence and violation of musical regularities—and that reward-related activations induced by music may be driven by the intrinsic value of successfully anticipating potential musical surprises (Koelsch et a., 2019; Gold et al., 2019; Cheung et al., 2019). Interestingly, humans show significant individual differences in sensitivity to musical pleasure, and this variance is related to brain structure and function in the reward circuitry, and its interactions with the auditory perceptual system (Mas-Herrero et al., 2013, 2014; Martinez-Molina et al., 2016, 2019).

In the current study, we explored the effects of dopamine on reward-potentiated music-memory encoding and consolidation. We aimed to evaluate whether music drives improvements in higher cognitive functions such as information encoding and long-term consolidation via the activation of the dopaminergic midbrain (e.g., SN/VTA-Hippocampal loop; Lisman et al., 2005, 2011; Adcock et al., 2006; Ripollés et al., 2016, 2018). In a recent *behavioral* study, we advanced this theoretical framework by collecting subjective ratings of pleasure from participants who listened to unfamiliar pieces of classical music. The results indicated that music-related pleasure and memory are, behaviorally, intimately related: the greater the pleasure elicited by a particular song, the better its ‘musical-memory recognition’ after a consolidation period (24 h), especially in individuals with high sensitivity to musical reward (Ferreri and Rodriguez-Fornells, 2017). Exploration of the neurochemical mechanisms underpinning this music-reward-driven effect on memory was not possible in this previous study, however, due to its behavioral focus. To that end, pharmacological interventions represent a promising and effective avenue for investigating the causal implications of dopamine-dependent mechanisms in learning and memory processes. Several studies, including our work on intrinsic reward from self-regulated learning (Ripollés et al., 2018), have shown that increasing synaptic dopamine concentration via d-amphetamine, methylphenidate (i.e., drugs blocking dopamine reuptake), or levodopa (i.e., a dopamine precursor) can enhance learning and memory performance in both healthy (Breinstein et al., 2004; Knecht et al., 2004; Whiting et al., 2007; 2008; Chowdury et al., 2012; Daniel et al., 2012; Bunzeck et al., 2014; Linsenn et al., 2014; Shellshear et al., 2015) and clinical populations (Berthier et al., 2011).

Given these antecedents, we aimed to investigate whether dopamine also plays a *causal* role in reward-potentiated-music *memory*. The question, in other words, is whether music induces reward-related responses will ultimately modulate long-term memory via dopaminergic transmission. We addressed these questions through a double-blind, within-subject pharmacological design in which we directly manipulated synaptic dopamine availability. Participants listened to unfamiliar music excerpts after orally ingesting a dopamine precursor (levodopa), a dopamine antagonist (risperidone), and a placebo across three sessions (same participants and methodology as in Ferreri et al., 2019 and Ripollés et al., 2018). In contrast to methylphenidate and d-amphetamines, levodopa does not indiscriminately enhance tonic dopamine levels but is rather rapidly taken up by dopaminergic neurons, transformed into dopamine, and stored in vesicles. Levodopa therefore increases the dopamine available for release each time a dopaminergic neuron fires. Risperidone interferes with dopaminergic neurotransmission by binding to and blocking D2-like dopamine receptors, which ultimately reduces the transmission of dopaminergic signals to post-synaptic neurons (Burstein et al., 2005).

The purpose of this study was to assess whether the modulation of the dopaminergic system influences music-related memory, rather than to explore the capacity of the drugs on their own to block or enhance the natural physiological responses induced by music. The dopaminergic system has an intrinsic physiological state that can be parametrized using the values measured in the course of a placebo session. Levodopa and risperidone were chosen specifically for their capacity to *displace* the dopaminergic physiological system from this baseline, and in opposite directions: risperidone reduces the effects of dopamine release, while levodopa enhances dopaminergic neurotransmission. As the aim was to bring the dopaminergic system *away* from its baseline state, our analyses directly compare the risperidone and levodopa data against each other by using the placebo session as a baseline (i.e., we present all data as the percentage of change from placebo, as in Ferreri et al., 2019 and Ripollés et al., 2018). Music-reward responses were measured by asking participants to provide subjective pleasure ratings after each musical excerpt. Ratings for arousal, emotional valence, and familiarity were collected as a control, among other measures. In order to test our main hypothesis, episodic memory performance for the presented songs was tested 24 hours after encoding using a recognition-recollection paradigm (Yonelinas, 2002).

For all planned analyses, we divided our population into two groups (high musical hedonia, HDN+; low musical hedonia, HDN-; as in Ripollés et al., 2018) according to their sensitivity to *musical reward* (i.e., their musical hedonia) using the Barcelona Music Reward Questionnaire (BMRQ, Mas-Herrero et al., 2013). The rationale for this split is twofold. First, reward-related behavioral and physiological responses to music are modulated by musical hedonia, as are the structure and function of core reward-related brain regions (Brattico and Pearce, 2013; Mas-Herrero et al., 2013, 2014; Martinez-Molina et al., 2016, 2019; Belfi et al., 2019; Belfi and Loui, 2019; Ferreri et al., 2019; Gold et al., 2019). Second, and most importantly, memory effects driven by dopamine in the context of intrinsic or abstract reward are highly dependent upon individual differences in reward sensitivity: the higher participants’ hedonia, the higher the memory and learning performance (Ripollés et al., 2018; Ferreri et al., 2017). Once participants were divided into two groups according to their BMRQ scores, mixed 2 × 2 repeated measures ANOVAs were calculated for each subjective rating and for memory recognition scores with Group as a between-subjects factor (HDN+, HDN-) and Intervention as a within-subjects factor (levodopa/risperidone, using the placebo session as a baseline). We predicted that, if reward-potentiated-music memory is a dopamine-dependent mechanism, these pharmacological interventions should modulate both musical pleasure and music memory performance for the more musical hedonic participants.

## Results

Analysis of the subjective ratings revealed that the pharmacological intervention modulated pleasantness (but not valence, familiarity, arousal, or *top ten* rankings, see methods) in interaction with the musical hedonia (significant Group × Intervention interaction: *F*(1,25)=4.479, *p*=.044, η^2^_p_ =0.152; no significant group or intervention effects, all p>0.272). This was further confirmed by paired Wilcoxon non-parametric tests (to better account for the N of each group of participants), which revealed that the difference in pleasantness ratings between levodopa and risperidone interventions (placebo-corrected values, see analysis section) was significant for the group with higher musical hedonia scores (*W* =17, p=.022) but not for the less hedonic participants (*W*=62, p=.263, see fig.1A). In other words: levodopa and risperidone increased and decreased, respectively, the pleasantness experienced by HDN+ participants while listening to unfamiliar musical excerpts. To unpack this interaction further, we calculated the general *drug effect* on pleasantness scores as the difference between the placebo-corrected effects induced by levodopa minus those induced by risperidone (same methodology I as in Ripollés et al., 2018 and Ferreri et al., 2019). This index allows us to test for a general effect of the pharmacological intervention (i.e., the effect of dopaminergic modulation) on a measure of interest. To test for a pleasantness drug effect between the HDN+ and HDN-groups, we used a non-parametric Mann-Whitney test, to better account for the number of participants in each group. As expected, the drug effect for pleasantness was significantly different between the groups (Mann-Whitney, *U*=41, p=.028, η^2^=0.227; Fig. 1A). A Spearman’s correlation confirmed that the higher the musical hedonia (i.e., BMRQ), the higher the drug effect on the pleasantness experienced (rs =.463, p=.030; fig.1C). No significant effects of pharmacological intervention, group, or interaction between the two were found for the other subjective ratings (arousal, emotional valence, familiarity, and *top ten* rankings, see methods; all *p* > 0.065).

**Fig. 1.**
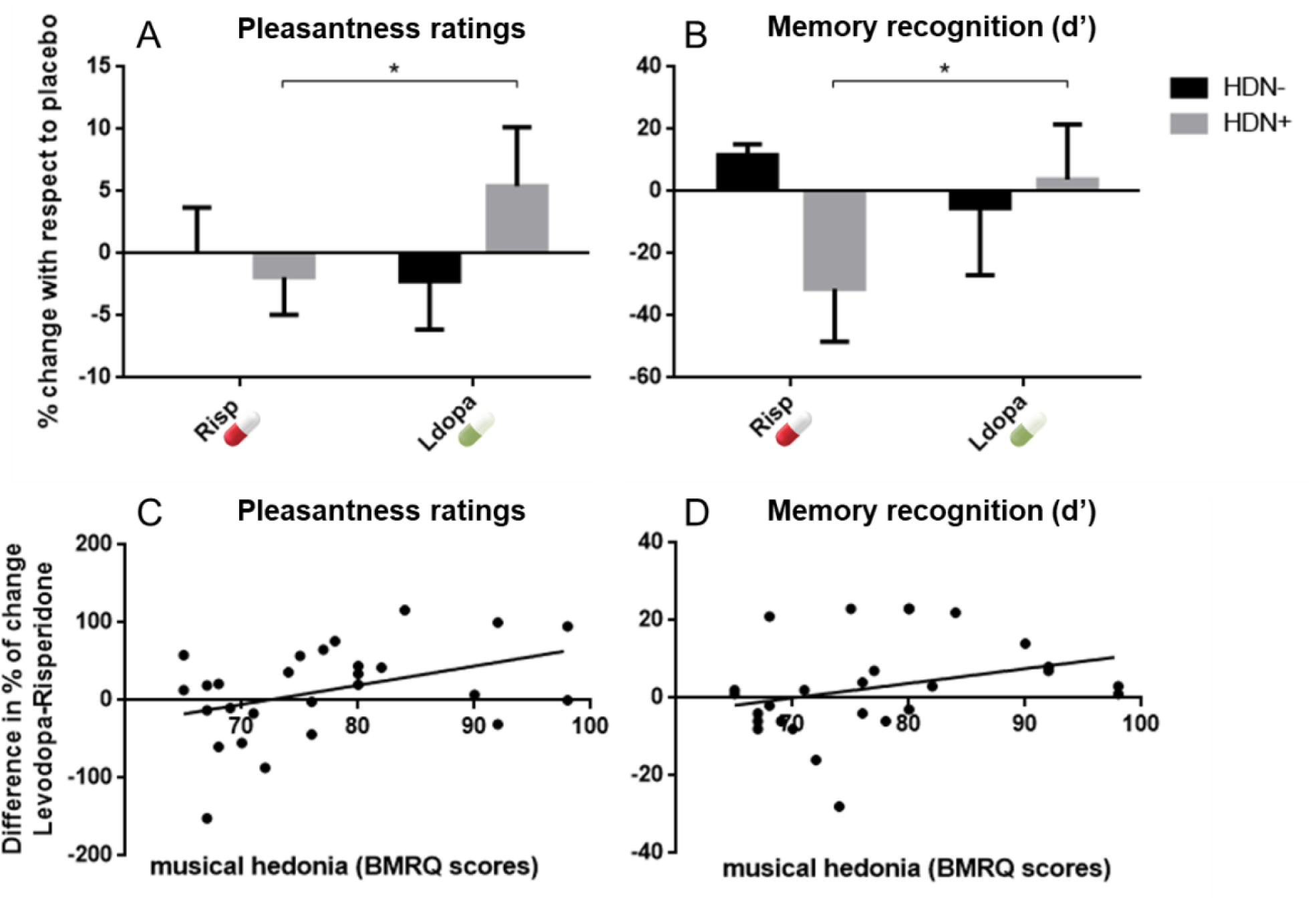
Effects of the pharmacological intervention on the pleasantness ratings (provided on Day 1) and music memory recognition (Day 2). Pleasantness ratings (**A, C**) and memory recognition (**B, D**) were calculated as the percentage of change from the placebo session (as in Ripollés et al., 2018 and Ferreri et al., 2019). Participants were divided according to their musical hedonia into HDN+ and HDN-groups using the median split of the BMRQ scores. Significant Group × Intervention interactions were obtained for both pleasantness (**A**) and memory recognition (**B**). In addition, the drug effect (i.e., the difference between the placebo-corrected levodopa and risperidone scores) was calculated for both pleasantness (**C**) and memory recognition (**D**) and was significantly correlated with individual differences in sensitivity to musical reward (the greater the musical hedonia, the greater the dopaminergic effect on pleasure and recognition memory). Note that, in the BMRQ larger values imply greater musical hedonia. Risp, risperidone. Levo, Levodopa, HDN+, High musical hedonia. HDN-, Low musical hedonia. Bars represent mean ± SEM.

Interestingly, the same pattern was found for music memory recognition (Fig. 1B). After 24 hours, participants completed an old/new recognition task, from which we computed d-prime scores (i.e., the discriminability index, obtained by dividing the difference between the means of the distributions for the old and the new items by the common standard deviation of the distributions; in other words: recognition effects) and corrected *remembered* (R) responses (i.e., number of R responses for old items minus number of R responses for new items; in other words: recollection effects) to assess recognition and recollection memory performance, respectively. The pharmacological intervention affected d-prime scores (i.e., they increased under levodopa and decreased under risperidone), for the HDN+ group (significant Group × Intervention interaction: *F*(1,25)=5.332, *p*=.029, η^2^_p_ =0.176; no other significant group or intervention effects, all *p*>.35). As expected, the difference in recognition memory between the two pharmacological interventions was significant for the HDN+ (*W*=18.5, p=.033) but not for the HDN-group (*W*=51.5; p=.701; see Fig. 1B). As with the pleasantness ratings, the general drug effect on memory performance was stronger for participants with higher musical hedonia scores (*U*=47, *p*=.033, η^2^=0.175). This result was further supported by a Spearman’s correlation showing that the higher the BMRQ scores, the larger the effect of the drug on recognition (*r*_*s*_=.400, *p*=.039; Fig. 1D). Thus, as with pleasure, the dopamine-dependent effect on music memory recognition was greater in participants with higher sensitivity to musical reward. No other significant effects for intervention, group, or the interaction between the two were found for *recollection* performance (all *p*>.286).

Finally, we found a positive correlation between the drug effects on pleasantness and on memory recognition: the more the drug effect modulated pleasantness ratings during musical listening, the greater the drug effect on music memory recognition was after the consolidation period (*r*_*s*_=.390, *p*=.044; Fig. 2). In other words, the stronger the *dopamine-dependent reward experience*, the larger the *dopamine-dependent memory effect*. No significant linear correlations or inverted U-shape relationships (Daniel et al., 2012) were found between either drug effect (on pleasantness or on recognition memory) and the weight-dependent drug dosage (all *ps*>.118).

**Fig. 2.**
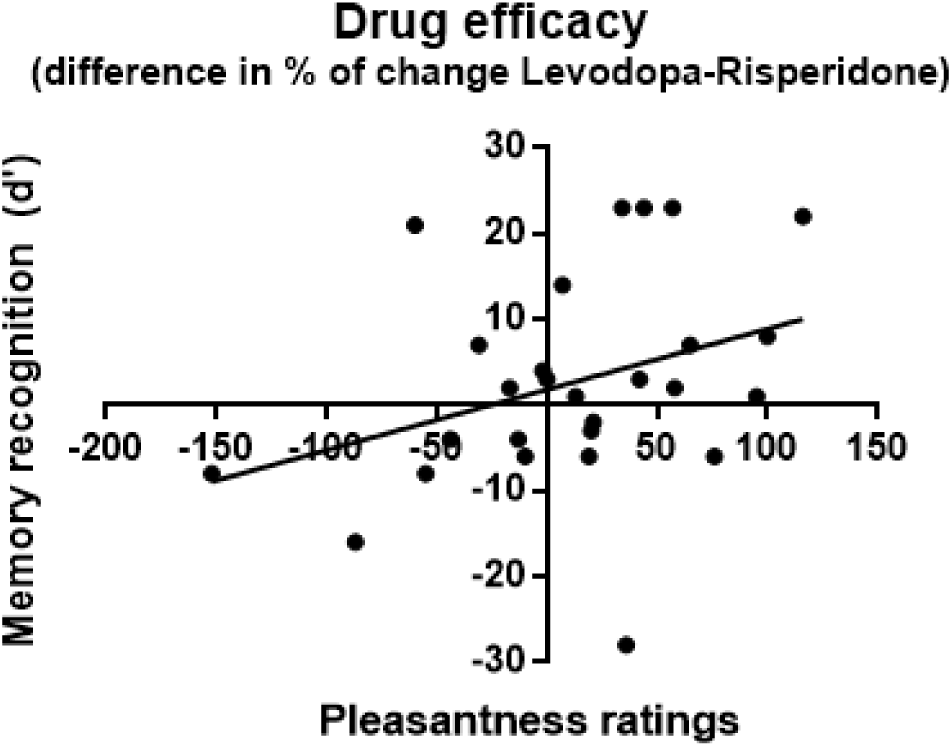
Relationship between the drug effects on pleasantness (Day 1) and memory recognition (Day 2), showing that the greater the drug modulation of pleasantness during music listening, the larger the drug effect on music memory recognition after a consolidation period.

## Discussion

Our findings show that dopaminergic synaptic availability, when manipulated through a within-subject, double-blind pharmacological paradigm, modulates both the pleasure and the recognition memory associated with one of the most iconic abstract rewards in humans: music. More specifically, we found that a dopamine antagonist (risperidone) decreased and a precursor (levodopa) increased, respectively, the level of pleasantness experienced by the participants during the encoding of unfamiliar musical excerpts, as well as their music memory recognition after a 24-hour consolidation period. However, our findings interestingly show (as in Ripollés et al., 2018) that the memory effects induced by the pharmacological intervention were only significant in participants with a higher sensitivity to musical reward (i.e., musically hedonic volunteers).

Previous research indicates a tight link between dopamine and musical pleasure. Key dopaminergic regions such as the ventral striatum (VS) and the midbrain respond to highly pleasurable musical stimuli (Blood and Zatorre, 2001; Salimpoor et al., 2011; 2013). Recent studies in which the dopaminergic reward system was manipulated via transcranial magnetic stimulation (Mas-Herrero et al., 2018) or through pharmacological interventions (Ferreri et al., 2019) suggest a *causal* role for the striatum and for dopamine, respectively, in both the ‘liking’ (i.e., pleasurable responses) and the ‘wanting’ (i.e., motivational responses) components of reward during music listening (Berridge et al., 2009). These studies did not, however, report individual differences in music-reward sensitivity: here we show that *dopamine-dependent* effects on pleasure and memory recognition vary with personal levels of musical hedonia.

In contrast to the previous studies on musical reward, in which pop, familiar, and favorite music were used as stimuli (Blood and Zatorre, 2001; Salimpoor et al., 2011), we used excerpts from unfamiliar pieces of classical music for our memory task. Given the high correlation between musical pleasure/motivation and familiarity (Salimpoor et al., 2011; 2013; Van Den Bosch et al., 2013), the unfamiliarity of the excerpts used here may have prevented some participants from experiencing intense pleasure and the associated boost in motivational responses. The purpose of this study was not, however, to assess the capacity of the drugs individually to block or enhance the natural physiological responses influenced by dopamine, but rather to elucidate whether modulation of the dopaminergic system influences music-related reward responses and memory. Interestingly, our results suggest that dopaminergic enhancement of musical pleasure and memory for non-familiar and non-preferred music (Ferreri et al., 2019) is observed only when participants can easily experience musical reward in their daily lives. In other words, dopamine cannot make you *enjoy* music more if you do not enjoy music that much in the first place. Importantly, individuals with low musical hedonia present a reduced connectivity between the VS and the auditory cortex (Martinez-Molina et al., 2015; 2019; Loui et al., 2018). These findings fit with a model according to which the engagement of the core-reward circuitry in response to music is triggered by signals from cortical auditory regions (Zatorre and Salimpoor, 2013). Thus, if the crosstalk between these two structures is not functional, no reward related activations occur while listening to music, independently of the integrity of the core reward circuitry (Martinez-Molina et al., 2015). In line with this interpretation, only individuals with high musical hedonia benefited from the intervention as reflected by a positive correlation between general drug effect on the subjective ratings of musical pleasure and the BMRQ scores. This result suggests that humans can selectively trigger intrinsic reward responses to behaviors or experiences to which they attribute special value, even in the absence of clear survival benefits or obvious links to any primary reinforcer (White, 1959; Berlyne, 1960; Gottlieb, 2013; Barto, 2013; Ripollés et al., 2016).

A main finding of the present study concerns the dopamine-dependent, reward-potentiated effect on music memory. Our results support previous research showing that rewarding stimuli enhance memory formation via dopaminergic pathways (Lisman and Grace, 2005; Adcock et al., 2006, Lisman et al., 2011; Wolosin et al., 2012; Ripollés et al., 2016). These findings are furthermore in line with previous pharmacological interventions showing that increasing the synaptic availability of dopamine enhances learning and long-term memory (Knecht et al., 2004; De Vries et al., 2010; Chowdhury et al., 2012; Shellshear et al., 2015; Ripollés et al., 2018). Crucially, and confirming the link between musical reward and musical memory, we showed that dopamine-dependent pleasurable responses provided during the encoding session were directly related to subsequent memory performance: the greater the effect of the drugs on pleasantness ratings, the larger the drug effect on recognition memory.

One possible interpretation of this finding relies on reward prediction mechanisms, which are widely known to increase dopamine release (Schultz, 1997; Watabe et al., 2017). Abstract rewards such as music are strongly dependent upon perceptual expectations and predictions (Meyer, 1956). In the context of prediction, data posits that dopaminergic neurons in the VS (and Nucleus Accumbens) are the key factor driving the attachment of hedonic value to music (Salimpoor et al., 2015; Gold et al., 2019; but see Cheung et al., 2019). Reward prediction errors (RPEs) are also crucial for reinforcement learning processes (Sutton and Barto, 2018), and growing evidence suggests that they play a pivotal role in episodic memory (Davidow et al., 2016; De Loof et al., 2018; Calderon et al., 2019). By experimentally manipulating RPEs during the encoding of faces, Calderon et al. (2019) recently showed that trial-specific RPE responses in the VS during learning predict the strength of the subsequent episodic memory. It is therefore possible that the dopamine-dependent RPEs underpinning musical pleasure during encoding might also promote episodic memory formation for the same material via the SN/VTA-Hippocampal loop (Davidow et al., 2016).

Of special note, and in agreement with our previous results that used the same pharmacological intervention to assay intrinsic reward as elicited from self-regulated learning (Ripollés et al., 2018), the current findings draw a complex picture of the relationship between abstract rewards and human memory: inter-individual differences in music reward sensitivity appear to play a crucial role in modulating not only the intense pleasure that music can evoke (Martinez-Molina et al., 2016), but also in musical memory formation. While in our previous work we showed that *learning itself* triggers intrinsic, dopamine-dependent reward signals that are amplified in more hedonic participants, here we show that *music itself* elicits dopamine-dependent reward signals that are comparably modulated by individual differences in musical hedonia. Indeed, our results suggest that music memory performance is driven not simply by synaptic dopamine availability, but also by the degree to which each individual can experience music reward in general (Ferreri and Rodriguez-Fornells, 2017). This in turn indicates new avenues for the study of the underlying mechanisms of music-driven memory benefits (Ferreri and Verga, 2016) and their implications in the clinical domain (e.g., Särkämo et al., 2008, 2014; Simmons-Stern et al., 2010; see Sihvonen et al., 2017 for review). By showing that musical reward is a crucial mechanism in music memory performance (Ferreri and Rodriguez-Fornells, 2017; but see also Grau-Sanchez et al., 2018; Särkämo, 2018 for the implication of musical reward in other domains), our results suggest that inter-individual differences in musical hedonia should be taken into account in memory stimulation and rehabilitation paradigms. Such broadened paradigms could facilitate the creation of more finely grained musical interventions in normal and pathological aging, for example (Ferreri et al., 2018). To that end, the current findings may represent an important first step in novel investigations of pathological aging since musical memory constitutes a special type of memory often spared also in disorders like Alzheimer’s disease (Baird and Samson, 2015). Further studies will help to elucidate whether a transfer effect of musical reward learning is also possible for the encoding of non-musical (e.g., verbal) material (Ferreri and Verga, 2016).

In conclusion, we show that pharmacologically increasing or decreasing dopaminergic signaling modulates behavioral measures of musical pleasure and long-term recognition memory for musical pieces, especially in individuals with high sensitivity to musical reward. This work, like our previous findings (Ripollés et al., 2016, 2018), emphasizes the versatility of the human dopaminergic reward system: dopamine signaling lies at the core of the memory benefits mediated not only by explicit or primary, but also intrinsic, abstract, and aesthetic rewards of different origins and modalities.

## Material and Methods

### Participants

Participant selection and procedure description have been previously illustrated in Ripollés et al., (2018) and Ferreri et al., (2019; participants are the same as in these two other studies). Around 150 individuals responded to advertisements and were contacted for a first phone prescreening. Of those, 45 confirmed their availability and were admitted at the hospital for further screening, medical examination and laboratory exams (blood and urinalysis). The study was approved by the Ethics Committee of Hospital de la Santa Creu i Sant Pau and the Spanish Medicines and Medical Devices Agency (EudraCT 2016-000801-35). The study was carried out in accordance with the Declaration of Helsinki and the ICH Good Clinical Practice Guidelines. All volunteers gave their written informed consent to participation prior to any procedure.

Subjects were judged healthy at screening 3 weeks before the first dose based on medical history, physical examination, vital signs, electrocardiogram, laboratory assessments, negative urine drug screens, and negative hepatitis B and C, and HIV serologies. The volunteers were excluded if they had used any prescription or over-the-counter medications in the 14 days before screening, if they had a medical history of alcohol and/or drug abuse, a consumption of more than 24 or 40 grams of alcohol per day for female and male, respectively, or if they smoked more than 10 cigarettes per day. Women with a positive pregnancy test or not using efficient contraception methods, participants with musical training, and those unable to understand the nature and consequences of the trial or the testing procedures involved were also excluded. Additionally, volunteers were requested to abstain from alcohol, tobacco, and caffeinated drinks for at least the 24 hours prior to each experimental period.

Twenty-nine volunteers completed the study (19 females, mean age=22.83±4.39) in exchange for a monetary compensation according to the Spanish Legislation. The original sample size was chosen to be 30 participants, but one participant dropped out early in the study and only 29 finalized it. This sample size was selected based on several criteria, including the recommendation that, in order to achieve 80% of power, at least 30 participants should be included in an experiment in which the expected effect size is medium to large (Cohen, 1988). In addition, we took into account the sample sizes of previous studies using levodopa to modulate memory (range: between 10 and 30 participants; Knecht et al., 2004; Copland et al., 2009; de Vries et al., 2010; Chowdhury et al., 2012; Apitz and Bunzeck, 2013; Shellshear et al., 2015). We also computed a sample size analysis using the G*Power program, which showed that a sample size of 28 was required to ensure 80% of power to detect a significant effect (0.25) in a repeated-measures ANOVA with three interventions at the 5% significance level. Selected participants were also tested with the extended version of the Barcelona Music Reward Questionnaire (BMRQ, Mas-Herrero et al., 2013), able to measure the individual sensitivity to musical reward (i.e., musical hedonia) and to explain individual differences in brain structure and function in response to pleasurable music (Mas-Herrero et al., 2013; 2014; Martinez Molina et al., 2016; 2019). We employed here an extended version of the BMRQ, including two items of the Montreal Battery of Evaluation of Amusia (MBEA, Peretz et al, 2003; see also Ferreri et al., 2019). Furthermore, participants were tested with the physical anhedonia scale (PAS, Chapman et al., 1976). No participants presented signs of amusia. Two participants scored within the ranges considered to indicate musical anhedonia and general anhedonia and were therefore excluded from the analysis here reported (total N=27, 18 females, mean age=23 ± 4.48, mean BMRQ=77.07 ± 9.89).

### Study design and procedure

This double-blind, crossover, treatment sequence-randomized study (Ripollés et al., 2018; Ferreri et al., 2019) was performed at the Neuropsychopharmacology Unit and Center for Drug Research (CIM) of the Santa Creu i Sant Pau Hospital of Barcelona (Spain). Experimental testing took place over three sessions (i.e. interventions). For each one, participants arrived at the hospital under fasting conditions and were given a light breakfast. Subsequently, they received in a double-blind masked fashion a capsule containing the treatment: a dopaminergic precursor with an inhibitor of peripheral dopamine metabolism (levodopa, 100 mg + carbidopa, 25 mg), a dopamine receptor antagonist (risperidone, 2 mg), or placebo (lactose). The dopaminergic system has a physiological or intrinsic state whose effects are most likely reflected by the values of the dependent variables measured during the placebo intervention. In this study, we intended to lower and raise this baseline dopaminergic state by means of two independent pharmacological interventions involving low-to-moderate doses of levodopa and risperidone. Drug doses were carefully chosen to be low enough to induce the desired modulation but not too large to allow collateral effects to become a confounding factor. In particular, the levodopa dose was kept in line with previous studies in healthy participants and within the dose range administered in clinical practice for the treatment of Parkinson’s disease. Drug doses use were decided upon these ethical concerns and the binding request on the part of our local Institute Review Board. After around 20 minutes of completing behavioral music tasks not described in the current manuscript (Ferreri et al., 2019), the participants completed a musical memory task which lasted 45 min approximately, followed by a language learning task (described in Ripollés et al., 2018). Next, participants spent their time in a resting room and were allowed to leave the hospital after 6 hr from the treatment administration. For each intervention, each participant came back 24 hours later for a behavioral memory retesting (without any pharmacological intervention), which lasted about 15 min. At least 1 week passed between one intervention and the other.

### Music memory task

This task has been validated and described in our previous work (Ferreri and Rodriguez-Fornells, 2017). In each intervention, participants were exposed three times to unfamiliar instrumental classical excerpts (normalized at − 10 dB, and faded 3 s in and 3 s out). During the first exposure, volunteers listened through earphones to 24 excerpts, lasting 20 s each (Eschrich et al. 2008. Participants were told to listen to the excerpts attentively, as they would be asked to remember them later. After each excerpt, they were asked to rate (1–5 points scales) the level of arousal, emotional valence, and familiarity, and the general pleasantness experienced when listening to the piece (i.e., “liking” reward measure, Berridge et al. 2009). Furthermore, we asked participants to indicate in which position of a top-ten classification (i.e. “wanting” reward measure, Berridge et al. 2009), they would like to place each excerpt, knowing that the excerpts ranked in the first three positions were more likely to be part of a final Spotify playlist that they were going to receive by e-mail for their participation. During the second exposure, participants were simply asked to listen again to the same excerpts, in order to be completely absorbed in music listening and encoding (Ferreri and Rodriguez-Fornells, 2017). During the third exposure, they were asked to listen to them another time and to rate again general pleasantness and top ten. The mean of the reward (i.e., pleasantness and top-ten) subjective ratings between the first and the third exposure were computed and employed for the analyses further reported (Ferreri and Rodriguez-Fornells, 2017). One minute passed between each exposure to all 24 musical excerpts.

24 hours after learning, participants were presented with 24 old and 24 new excerpts, lasting 10 s each. The selection of these 10 s pieces (Halpern and Müllensiefen 2008) was made by excluding the first and last 3 s (i.e., the faded ones) of the excerpts and by selecting at least one musical phrase. For each one, participants had to indicate if they listened to it the day before (old/ new recognition). If they thought they had, they neeeded to commit to one of three additional options (recollection task): remember (R), know (K), or guess (G). R indicated that they could recollect something specific about the study episode; K indicated that the excerpt was confidently familiar, but they had no recollective experience; G responses were given when unsure about whether the excerpt was really heard the day before(R/K paradigm; Yonelinas, 2002). In total, six lists of excerpts (balanced for emotional valence, arousal, general pleasure and familiarity) were presented to each participant: three lists (one for each intervention) during the encoding session (i.e., old) and three during the test session, 24 hours later (i.e., new). The order of the lists was counterbalanced across interventions. The six lists were created (pretested on N=60 participants, 44 females, mean age = 28 ± 12.08) so that there were no differences (one-way ANOVA and Bayes Factors) in arousal (*F*(5,115)=0.061; p=0.997; η^2^ =0.003; BF_10_=0.019), emotional valence (*F*(5,115)=0.193; p=965; η^2^ =0.008; BF_10_=0.024), general pleasure (*F*(5,115)=0.325; p=0.897; η^2^ =0.014; BF_10_=0.033; BF_10_=0.031), and familiarity (*F*(5,115)=0.371; p=0.868; η^2^ =0.016; BF_10_=0.033). Items that were judged by participants as familiar or very familiar (with a rating >=4) in the current experiment were excluded from the analyses here reported. The total duration of this retrieval phase lasted about 20 min.

Auditory stimuli were presented using a headset, and the overall loudness of the excerpts was adjusted subjectively to ensure constant loudness throughout the experiment.

### Analysis

In this study, the main aim was to bring the dopaminergic system away from its intrinsic state (i.e., the placebo intervention) and in opposite directions. Risperidone and levodopa were chosen to lower and increase the dopaminergic basal state of the participants, respectively. Thus, for both subjective ratings (day 1, e.g., pleasantness, top-ten, arousal, emotional valence and familiarity) and memory performance (day 2), we computed the percentage of change under risperidone and levodopa with respect to the placebo section (Ripollés et al., 2018; Ferreri et al., 2019). D prime scores (i.e., the discriminability index, obtained by dividing the difference between the means of the distributions for the old and the new items by the common standard deviation of the distributions) and corrected R responses (i.e., number of R responses for old items minus number of R responses for new items) were computed for testing memory scores, namely recognition and recollection memory performance, respectively. In order to test the implication of musical reward sensitivity in a drug-dependent reward effect on memory (Ferreri and Rodriguez-Fornells, 2017), we computed the median BMRQ value to split our final sample of 27 participants into high and low musical hedonic groups (Ferreri and Rodriguez Fornells, 2017; Ripollés et al., 2018). We therefore run a mixed repeated measures ANOVAs for each measure (i.e., subjective ratings and memory scores) with drug intervention (i.e., placebo-corrected percentages of change) as within-subject, and BMRQ group (i.e., low or high musical hedonia) as a between-subject factor.

To unpack the results of the mixed repeated measures ANOVA, we used paired Wilcoxon non parametric tests to compare the placebo-corrected values under risperidone vs levodopa in the two groups of volunteers. We then calculated the general drug effect for significant interactions between interventions (risperidone, levodopa) and group (musical hedonia). This was calculated as a subtraction of the percentage of change from placebo induced by levodopa minus the percentage of change from placebo induced by risperidone intervention (see, for a similar analysis, Ripollés et al., 2018). This measure allows us to test for a general effect of the pharmacological intervention (i.e., the effect of dopaminergic modulation) on each measure of interest. We then assessed if the general drug effect for the measures showing a significant interaction were different for high vs. low musical hedonic groups, by using a non-parametric Mann-Whitney test to better account for the reduced number of participants for each group. Additionally, we used Spearman’s correlations to analyse (in a non-discrete manner) the relationship between musical reward sensitivity (individual BMRQ scores) and the general drug effect for the measures showing a significant interaction. These between group tests and correlations are FDR corrected with a p<0.05 threshold.

Furthermore, in order to test the relationship between the drug-dependent modulations of subjective ratings and recognition memory, we performed a follow up analysis in which we correlated the drug effects of the measures for which we previously found significant interactions (i.e., pleasantness and memory recognition). Finally, as a control and based on previous research (Chowdury et al., 2012), we correlated (Spearman’s correlations) the drug effect of significant interactions with a weight-dependent measure of drug dose, calculated in mg of levodopa/risperidone per kilogram. For significant interactions of mixed between-within ANOVA models, partial eta squares (η^2^_p_) are provided as measures of effect size. For significant differences measured with the Mann-Whitney or Wilcoxon test, eta squares (η^2^) are provided (calculated as Z^2^/N-1). In addition, to ensure that the lists of songs were equal in terms of familiarity, aorusal and valence, pleasantness and familiarity, confirmatory Bayesian statistical analyses were computed with the software JASP using default priors (JASP Team, 2018; Morey et al., 2015; Rouder and Morey, 2012). We reported Bayes factors (BF_10_), which reflect how likely data is to arise from one model, compared, in our case, to the null model (i.e. the probability of the data given H1 relative to H0).

## ACKNOWLEDGMENTS

We thank the staff of the Centre d’Investigació del Medicament de l’Institut de Recerca HSCSP for their help. The present project has been funded by the Spanish Government (MINECO Grant PSI2011-29219 to A.R.F.) L.F. was partially supported by Morelli-Rotary postdoctoral fellowship and IMPULSION (IDEX Lyon) grant. M.V. was partially supported by FIS trough grant CP04/00 121 from the Spanish Health Ministry in collaboration with Institut de Recerca de l’Hospital de la Santa Creu i Sant Pau, Barcelona; she is a member of CIBERSAM (funded by the Spanish Health Ministry, Instituto de Salud Carlos III). We thank Michael McPhee for his thoughtful comments and efforts towards improving this manuscript.

## ADDITIONAL INFORMATION

### Ethics

This study was performed according to local ethics and to the Declaration of Helsinki. It was approved by the Ethics Committee of Hospital Sant Pau and by the Spanish Medicines and Medical Devices Agency (EudraCT 2016-000801-35). All participants gave informed written consent and received compensation for their participation in the study according to Spanish legislation.

